# Single-molecule diffusometry reveals the nucleotide-dependent oligomerization pathways of *Nicotiana tabacum* Rubisco activase

**DOI:** 10.1101/191742

**Authors:** Quan Wang, Andrew J. Serban, Rebekka M. Wachter, W.E. Moerner

## Abstract

Oligomerization plays an important role in the function of many proteins, but a quantitative picture of the oligomer distribution has been difficult to obtain using existing techniques. Here we describe a method that combines sub-stoichiometric labeling and recently-developed single-molecule diffusometry to measure the size distribution of oligomers under equilibrium conditions in solution, one molecule at a time. We use this technique to characterize the oligomerization behavior of *Nicotiana tabacum* (Nt) rubisco activase (Nt-Rca), a chaperone-like, AAA-plus ATPase essential in regulating carbon fixation during photosynthesis. We directly observed monomers, dimers and a tetramer/hexamer mixture, and extracted their fractional abundance as a function of protein concentration. We show that the oligomerization pathway of Nt-Rca is nucleotide dependent: ATPγS binding strongly promotes tetramer/hexamer formation from dimers and results in a preferred tetramer/hexamer population for concentrations in the 1-10μM range. Furthermore, we directly observed dynamic assembly and disassembly processes of single complexes in real time, and from there estimated the rate of subunit exchange to be ~0.1s^-1^ with ATPγS. On the other hand, ADP binding destabilizes Rca complexes by enhancing the rate of subunit exchange by >2 fold. These observations provide a quantitative starting point to elucidate the structure-function relations of Nt-Rca complexes. We envision the method to fill a critical gap in defining and quantifying protein assembly pathways in the small-oligomer regime.

## Introduction

The oligomerization state of many proteins is critical to their function^1^. It forms the basis for substrate recognition for many enzymes^2^ and underpins the rich biophysical phenomena of allostery^3^ and cooperativity^4^,^5^. On the other hand, erroneous aggregation is involved in the pathogenesis of various amyloid diseases (e.g. Alzheimer’s, Parkinson’s, Huntington’s, etc.)^6^,^7^. Understanding the mechanism and pathways by which proteins assemble into higher order structures is of fundamental and biomedical significance. Several biochemical and biophysical methods are available to examine the oligomerization behavior of proteins but all have certain limitations. Methods based on liquid chromatography and sedimentation result in at least partial fractionation of the sample, an effect that is not suitable for complexes that undergo rapid exchange during measurements. Scattering-based techniques like small angle X-ray scattering (SAXS)^8^ and dynamic light scattering (DLS) rely on model-dependent fitting, which can be error-prone for highly polydispersed and/or interacting samples. Recently, native mass spectrometry has shown great promise in characterizing the quaternary structure of proteins, but the degree of perturbation introduced by electrospraying macromolecules into the gas phase is still debated^9^.

On the other hand, useful methods with single-molecule sensitivity have been developed. Electron microscopy offers structural information at the single-particle level but requires samples to be moved out of equilibrium onto a surface and dried in a vacuum. Recently, fluorescence correlation spectroscopy has been applied to resolve oligomerization states of proteins^10^,^11^ in solution, but this method has limited resolution for a mixture of assembly states^12^. Other techniques, such as two-color coincidence detection^13^, stepwise photobleaching^14^ and single-molecule FRET have been applied to characterize oligomerization under certain assumptions. Here, we describe a general method to directly measure the oligomer distribution of proteins in solution, with single-molecule resolution under equilibrium conditions. The method, termed “ABEL-oligo”, is based on measuring the hydrodynamic radius of individual molecular oligomers contained in an aqueous-phase Anti-Brownian ELectrokinetic (ABEL) trap in real-time. We use this method to dissect the oligomerization pathway of tobacco Rubisco activase, a catalytic chaperone essential for carbon fixation.

Rubisco activase (Rca) is a chaperone-like, AAA+ ATPase that rescues the main carbon fixation enzyme Rubisco [ribulose-1,5-bisphosphate (RuBP) carboxylase/oxygenase] from premature inhibition^15^–^17^. Despite its importance, the detailed mechanism of Rubisco activation by Rca is poorly understood, in part due to the complicated oligomerization behavior of Rca. Previous biophysical characterization of several higher plant Rca proteins using a suite of established methods^18^ revealed a highly heterogeneous and transient oligomer distribution, but these approaches could not be used to establish a quantitative oligomerization pathway. Surprisingly, sedimentation velocity experiments on tobacco (*Nicotiniana tabacum*, Nt) Rca^19^,^20^ suggested that the oligomerization of Nt-Rca does not depend on nucleotide. On the contrary, the assembly of spinach Rca was shown to be strongly dependent on the type of nucleotide^20^, consistent with earlier work reporting that both ATP and ATPγS promote the formation of higher-order oligomers (MW > 550kDa), whereas ADP does not^21^. Additionally, the oligomerization behavior of cotton β-Rca, as monitored by FCS, appeared to be a function of nucleotide.^22^. Meanwhile, several structural studies suggested hexamers as the functional species^17^,^23^,^24^ (Figure 1B). Here we set out to apply our ABEL-oligo method to define and characterize the assembly pathway of Nt-Rca *in vitro* and examine the effects of ATP binding (with Mg·ATPγS) and hydrolysis (with Mg·ADP).

**Figure 1.**
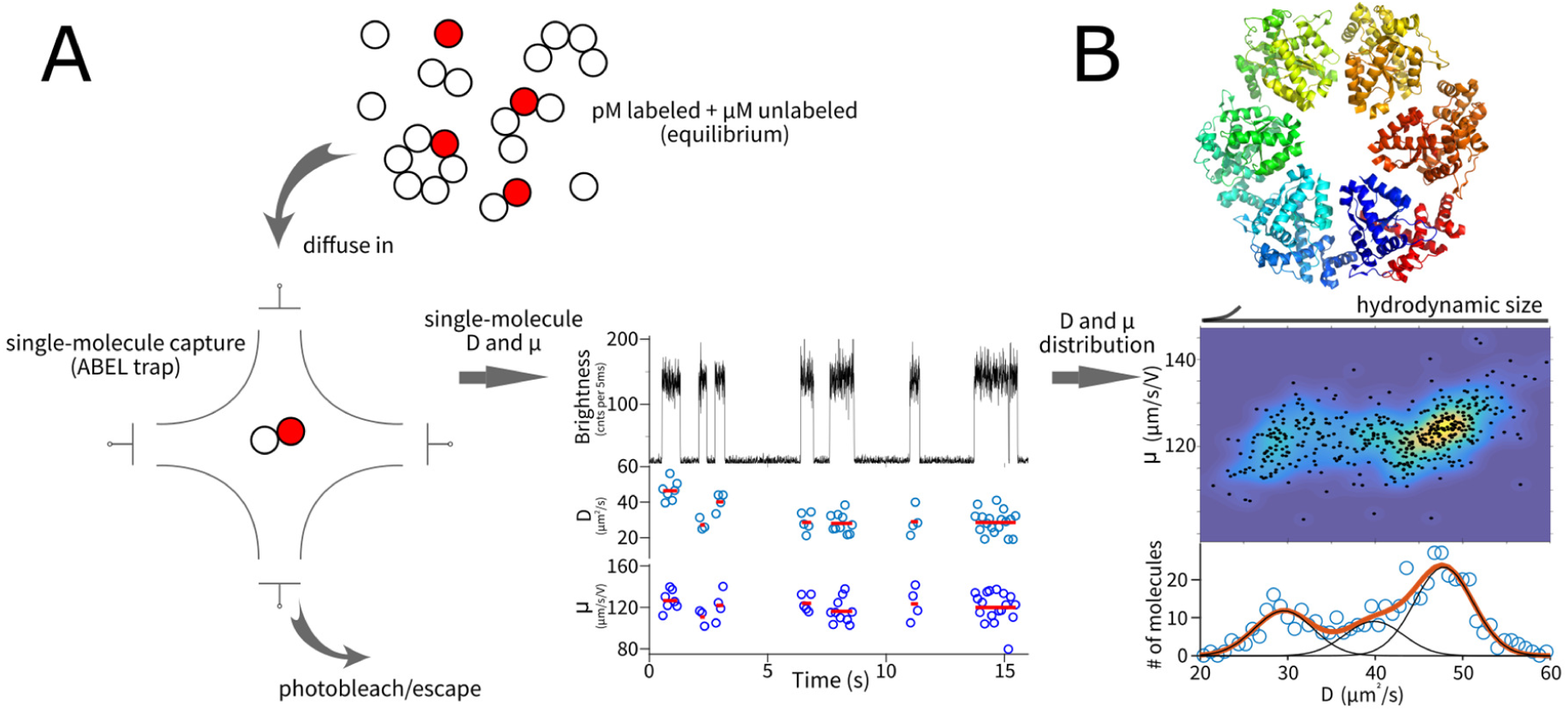
Measurement scheme. (A) Workflow of the assay based on single-molecule diffusometry. See main text for details. (B) Structural model of the closed-ring hexamer of truncated Nt-Rca (PDB: 3ZW6). Full-length protein chains were measured in this work.

## Materials and Methods

### Site-Directed Mutagenesis, Protein Expression and Purification

The gene coding for *Nicotiana tabacum* (tobacco, Nt) Rca (383 amino acid residues, MW 42.804 kDa, theoretical PI 5.5) was transferred into the pHUE plasmid^25^,^26^ following published procedures^27^. The QuikChange mutagenesis kit (Agilent Genomics) was used to introduce the S379C substitution near the C-terminus of Nt-Rca, as confirmed by DNA sequencing. Nt-Rca-S379C was expressed in *E. coli* BL21*(DE3) (Invitrogen) as a 6His-tagged ubiquitin fusion protein, and purified via affinity chromatography on a Ni^2+^-nitrilotriacetic acid (Ni-NTA) column (Qiagen). Subsequently, the N-terminal 6His-ubiquitin tag was cleaved using a deubiquitylating enzyme, the sample was dialyzed, and reapplied to a Ni-NTA column, as described previously^27^. Protein fractions were spin-concentrated and buffer-exchanged into 25 mM HEPES pH 7.5, 250 mM KCl, 5 mM MgCl_2_, 10% glycerol and 2 mM ADP. Protein preparations were quantitated using the Bradford method with bovine serum albumin as a standard. Aliquots were flash-frozen in liquid N_2_ and stored at - 80°C.

### Protein labeling method

The conjugation of the AlexaFluor-647 C2-maleimide fluorophore (Molecular Probes, Eugene, OR) to the engineered cysteine residue of Nt-Rca-S379C was carried out as described previously^22^,^27^. Briefly, labeling was carried out in the presence of 5 mM ADP at a dye:protein ratio of 3.5 in HEPES buffer pH 7.2. In control reactions containing wild-type Nt-Rca instead of Nt-Rca-S379C, the covalent labeling of protein was not observed, indicating that the cysteine residue in position 379 is specifically targeted by the maleimide functional group. Alexa-labeled Nt-β-Rca-S379C was purified by gel filtration, then analyzed by reverse-phase HPLC (Agilent Technologies 1100) on a C18 analytical column (Vydac 218 TP) using a linear water/acetonitrile gradient with 0.1% trifluoroacetic acid (TFA). Elution of protein and dye were monitored by optical density (OD) at 220, 280, and 650 nm. Protein eluted at 46.9 % acetonitrile as a single peak and carried the Alexa-dye label, whereas free Alexa-dye was not detected (expected elution at 16.4 % acetonitrile). Alexa/protein ratios were determined by absorbance measurements on collected HPLC fractions as described previously, and provided a 1:1 molar ratio of protein to dye^10^. Aliquots of labeled (88.2 μM) and label-free (268.0 μM) Nt-Rca-S379C in buffer containing 25 mM HEPES-NaOH, pH 7.5, 250 mM KCl, 2 mM ADP, 5 mM MgCl2, 10% glycerol were flash frozen in liquid N_2_ and stored at -80°C.

### Sample preparation for ABEL-oligo measurements

On the day of the experiment, samples were thawed on ice. Labeled Rca was first diluted to 3nM concentration in a buffer containing no Mg^2+^ or nucleotide (50mM HEPES, pH 7.5, 150mM KCl) and kept on ice for ~20min. Subsequently, a reaction mixture containing ~10pM labeled Rca and an appropriate amount of unlabeled Rca was created in the reaction buffer containing Mg^2+^ and nucleotide (50mM HEPES, 150mM KCl, 3mM Trolox, 0 or 5mM MgCl_2_, 0 or 2mM nucleotide, 10% glycerol). The mixture was incubated on ice for ~20min, deoxygenated (see below) and injected into the ABEL trap for single-molecule measurements. For cotton Rca, the 20min incubation time has been shown to be sufficient to reach equilibrium^10^. The concentration of labeled Rca was kept at ~10pM in all measurements. In experiments with 2mM ATPγS, the final solution also contained between 0 and 80μM ADP, which was carried over from the storage buffer of the unlabeled Rca preparation. Likewise, the low-ADP condition contained between 0 and 80μM ADP.

### ABEL trap setup

The ABEL trap was implemented as previously described^28^. Briefly, the apparatus consists of a position-sensing module, a signal-processing unit and a microfluidic sample holder. To sense the position of a single labeled biomolecule or oligomer in real time, we used fast laser scanning in a deterministic pattern and photon-by-photon molecule localization defined by the position of the laser spot at the moment of detection of a fluorescence photon. The signal-processing unit computes the most-probable position of the diffusing molecule given all available physical information. It is implemented on a field programmable gate array (FPGA) board for minimum computational delay. Given estimated position values, feedback voltages were generated, amplified and applied to the solution in a microfluidic sample holder to compensate for the Brownian motion of a single molecule. Our ABEL trap suppresses the Brownian motion of a single molecule in 2D (xy) only; diffusion in z is confined by the axial thickness of the microfluidic cell to ~700nm.

### Single-molecule diffusometry

The diffusion coefficient and electrokinetic mobility of individual trapped molecules were extracted using a maximum-likelihood estimator as previously described^29^. This works even though the molecule is prevented from diffusing away, because at any given time, the molecule is still driven by thermal diffusion as well as by the fields used for trapping. Briefly, the algorithm uses the photon-stamped position sequence of a single trapped molecule and the feedback voltages as inputs and utilizes an iterative procedure to reconstruct the motion trajectory. It then separates the in-trap motions of a single molecule into a voltage-dependent component and a stochastic component to estimate electrokinetic mobility and diffusion coefficient, respectively. The algorithm can be configured to run in real-time, or with higher precision offline, with a tunable time resolution between 50ms to ~1s. In general, both single-molecule size (from diffusion coefficient) and charge (from electrokinetic mobility) information can be extracted, as previously demonstrated for DNA hybridization^29^. In this work, the electrokinetic mobility is dominated by the electroosmotic response of the bulk solution (resulting from the polyelectrolyte multi-layer coating (see below) to prevent protein sticking) and contains little information about protein charge. We thus utilized only the diffusion information to identify the different oligomers.

### Conditions for single-molecule fluorescence detection

We used the 633nm line from a HeNe laser as the excitation source with an average excitation intensity of 1.5 kW/cm^2^. For experiments that aimed to probe the dynamic assembly/disassembly processes, the intensity was lowered to 0.75 kW/cm^2^. Trapping experiments were performed in the reaction buffer (50mM HEPES, pH 7.5, 150mM KCl, 3mM Trolox, 0 or 5mM MgCl_2_, 0 or 2mM nucleotide, 10% glycerol) with an added oxygen scavenging system^30^ (2mM protocatechuic acid and 50nM protocatechuate 3,4-dioxygenase) to prevent premature photobleaching of the dye Alexa647. To prevent protein sticking to the microfluidic surfaces, the sample chamber was passivated by polyelectrolyte multilayers using layer-by-layer deposition^31^,^32^. Sticking is easily detected by observation of the appearance of non-random behavior in the trapping field direction.

### Data Analysis

In the experiment, we only probe protein complexes containing at least one fluorescent subunit. We need to establish a link between the measurable quantities and the underlying oligomer distribution. The probability of an n-mer incorporating exactly one labeled subunit (*L*=1) assuming independence is given by the binomial distribution,

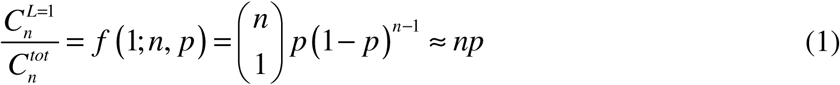

where the denominator is the total concentration of an n-mer, and *p* is the ratio between labeled and total Rca subunits in solution. Labeled Rca was kept at ~10pM and the total unlabeled Rca subunit concentration was varied between 200nM and 11μM. Under these conditions, *p* lies between 10^-6^ and 10^-4^, so (1-*p*)^n-1^ γ1 and oligomers with more than one labeled subunit can be ignored.

We can now relate the fractional concentration (P^n^) of the different oligomers to the apparent percentage concentrations measured using the fluorescent subset.

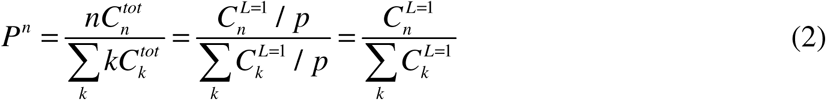

From the above equation, it becomes clear that by measuring the fluorescent subset of particles and the oligomeric stoichiometries n, we can extract the fractional concentrations of the different oligomers.

The changes in diffusion coefficient were detected by a change-point algorithm similar to Ref.^33^ but modified to use a Gaussian-based log-likelihood ratio test instead of a Poisson model as in the original publication. Two-dimensional densities were estimated from a scatter plot using a Gaussian-smoothing algorithm (2D kernel density estimation^34^).

For a monomer-dimer-tetramer-hexamer assembly model, we have

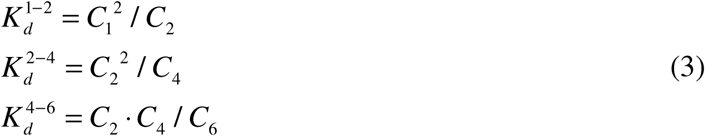

The *K*_*d*_ values were determined by a least-square procedure minimizing the following error function

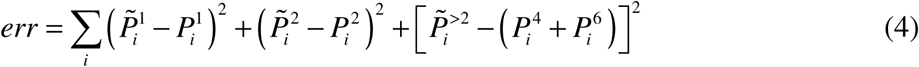

where 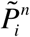 is the experimentally extracted fractional concentration of a n-mer at the i-th concentration point and 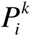 is the fractional concentration predicted by the equilibrium (Eq. 3) at the corresponding protomer concentration, 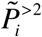 represents the unresolved tetramer/hexamer mixture seen in the experiment. This procedure can reliably extract 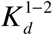 and 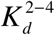 but not 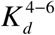 (the error function is insensitive to the value of 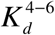). As a result, only 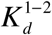 and 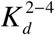 are reported in this work.

## Results

### Resolving oligomers under equilibrium conditions using single-molecule diffusometry

To resolve the different oligomerization states of Nt-Rca in solution, we first incubated labeled Nt-Rca (~10pM) with unlabeled proteins (0-11μM of protomer concentration). After equilibrium was reached, labeled Nt-Rca was randomly incorporated into the different assembly states. We then injected the equilibrated sample into a microfluidic-based single-molecule trapping system known as the Anti-Brownian ELectrokinetic (ABEL) trap described in Materials and Methods (Figure 1A). Inside the trap, individual fluorescently-tagged proteins were detected in solution with their diffusive motion greatly suppressed, resulting in long-time monitoring of single-molecule behavior in aqueous buffer solution. In the trap, individual proteins enter the center (“trapping region”) of the microfluidic chamber by diffusion and, if they carry a fluorescent subunit, are detected and captured by the feedback electrokinetic forces. These trapping events manifested themselves as brightness plateaus lasting ~1 second (Figure 1A). Termination of a trapping event is most likely due to photobleaching of the fluorescent label (Alexa647). When a single protein was captured, its diffusion coefficient and electrokinetic mobility were estimated with ~100ms time resolution using a statistical learning procedure^29^. The diffusion coefficient (*D*) is related to the hydrodynamic radius (*r*) of the molecule through the Stokes-Einstein relation (*D=k*_*B*_*T/6πηr*), and directly probes oligomerization state. Here, the electrokinetic mobility (*μ*) is only weakly dependent on molecular charge, because trapping was dominated by electro-osmotic forces resulting from charged, anti-fouling surface coatings of the microfluidic chamber (Materials and Methods). As a result, mobility values were expected to be similar for all measured molecules, regardless of their physical properties.

The ability to measure single-molecule size allowed us to probe the full distribution of oligomerization states in solution. We thus averaged all *D* values and all *μ* values collected for each individual protein complex. For all measured molecules, the average values were displayed as single points on a *D-μ* parameter space and also shown as a density distribution plot using a kernel density estimation^34^ method. In the example data shown in Figure 1A, obtained with 400nM Nt-Rca in the presence of ATPγS and Mg^2+^, we found three components in equilibrium with each other, as quantified by multi-Gaussian fitting of the projected *D* histogram. Sub-stoichiometric labeling and single-molecule diffusometry thus provide a powerful means to dissect the distribution of oligomerization states in solution under equilibrium conditions.

### The dimer is an Nt-Rca assembly intermediate

We conducted single-molecule diffusometry measurements with varying protein (subunit) concentrations [Rca] from 0-11μM, under three different nucleotide conditions. The results are summarized in Figure 2. Below 200nM, we saw exclusively a single population with a diffusion coefficient of ~50 μm^2^/s, which we identified to be the monomeric species. This is consistent with ultracentrifugation and SAXS data on Nt-Rca^20^. In further analysis, we thus normalized the diffusion coefficient of each complex by that of the monomer. At intermediate protein concentrations between 0.4μM and 2μM with no Mg^2+^ and low ADP (Figure 2 left), we observed multi-modal distributions, indicating assembly into well-defined oligomeric states. When fitting the projected D histogram (Figure 1A) using 3 Gaussians, we consistently resolved a distinct population with a D that is 80% of the monomer. Based on the simple D_n_/D_1_ = (1/n)^1^/^3^ (0.79 for n=2) scaling, we identified the dimer as an assembly intermediate under the low ADP condition. We note that although this simple scaling law is derived by approximating proteins as spheres, a consideration of more realistic protein shapes or structural models has been demonstrated to result in only minor corrections (D_2_/D_1_ from 0.79-0.80)^10^. Similarly, following the same analysis, dimers were also identified as assembly intermediates under Mg·ATPγS (Figure 2 middle) and Mg·ADP (Figure 2 right) conditions.

**Figure 2.**
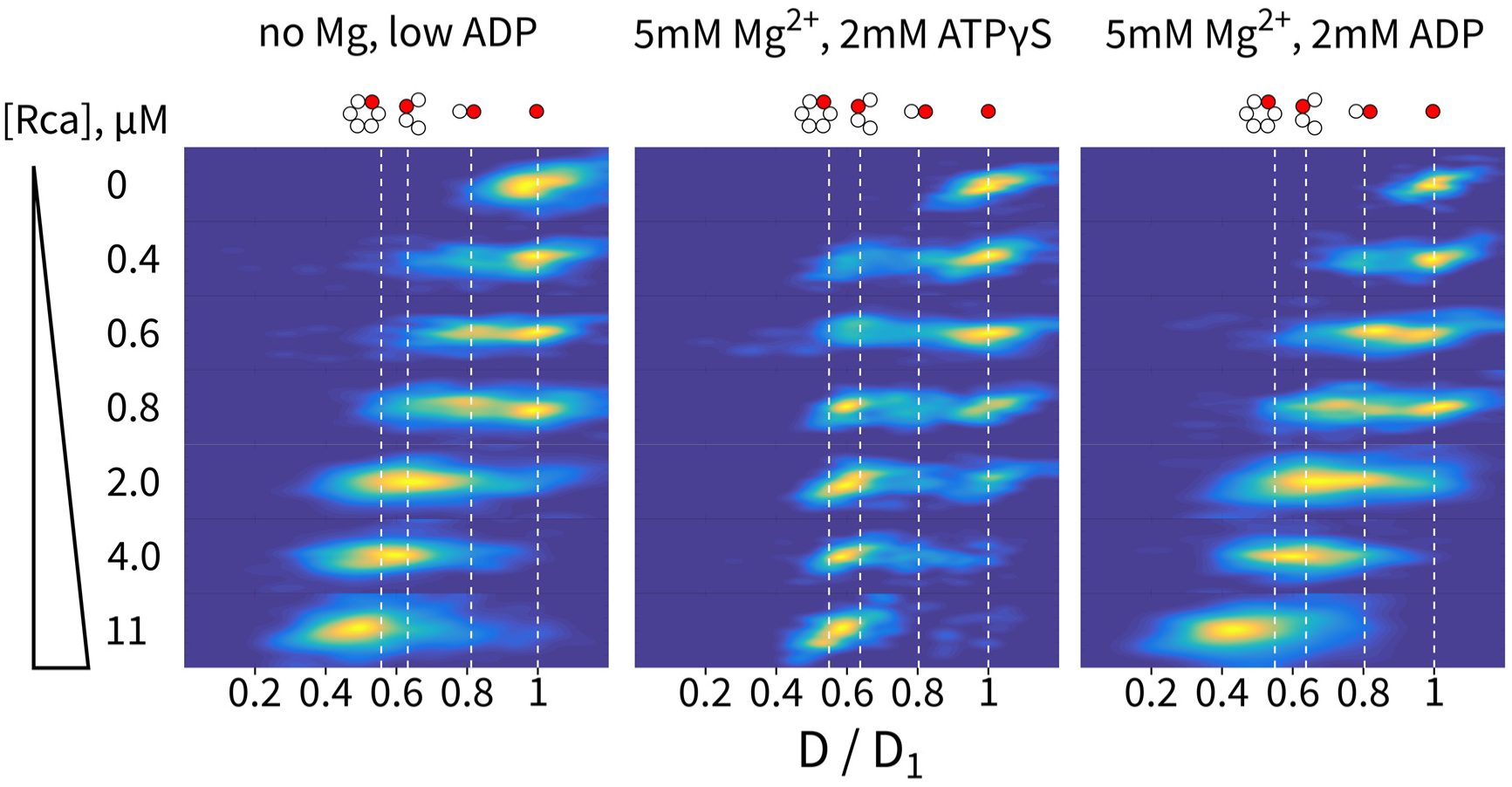
Mapping the distribution of Nt-Rca oligomers under different nucleotide binding conditions at varying protein concentrations. Density data are displayed with a normalized color scale.

### Oligomerization of Nt-Rca is nucleotide-dependent

Upon visual inspection, the oligomerization pathways are distinct for the different nucleotide conditions (Figure 2). To understand these differences more quantitatively, we highlight the expected D positions of the monomer, dimer, tetramer and hexamer species on the D-μ map using the D_n_/D_1_ = (1/n)^1^/^3^ scaling law. In the first condition (Figure 2 left), Mg^2+^ was omitted and residual ADP concentrations ranged from 0 to 80 μM. Since the K_i_ value for ADP inhibition of Nt Rca has recently been determined to be 37 μM^35^, we estimate that on average, about half of the active sites were occupied with nucleotide. Under these conditions, we observed the buildup of dimers and the appearance of tetramers in the range of 0.4 to 0.8μM Rca. At 2μM, Nt-Rca seemed to exist in a complex equilibrium containing monomers, dimers, tetramers and hexamers. Above 4μM, we observed near complete depletion of the monomer species while most of the population was found to exist as a mixture of tetramers and hexamers. At 11μM, oligomers larger than hexamers were observed. On the other hand, in the presence of 5 mM Mg^2+^ and 2 mM ATPγS, we observed a dramatically different oligomerization progression. Between 0.4 and 0.8μM, we observed weak dimer buildup and rapid formation of a D~0.6D_1_ species. We identified this state to be a mixture of tetramers and hexamers, given the limited resolution in measuring D values^29^ compared to the relatively small difference in D between tetramers and hexamers (~14% difference). Between 2 and 11μM, we observed a gradual sharpening of the tetramer/hexamer population. At 11μM, Nt-Rca seemed to be “locked” into the tetramer/hexamer form, instead of continuing to form higher oligomers, as observed for the low-ADP condition (Figure 2 left). We then conducted measurements in the presence of 5 mM Mg^2+^ and 2 mM ADP, which may mimic the post-hydrolysis state of the enzyme complex. In this case, we observed an oligomerization progression very similar to that of the low-ADP protein. Dimers populate at sub-μM concentrations, tetramers and hexamers exist between 1-4μM and aggregates larger than hexamers form at >10μM.

To further quantify the different oligomerization behavior as a function of nucleotide binding, we extracted the fractional concentrations of the different oligomers, defined to be 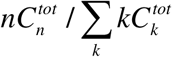 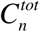 is the total concentration of an n-mer), directly from the data and fit them to an assembly model defined by the monomer-dimer-tetramer-hexamer pathway. Due to our limited ability to separate tetramers and hexamers, we combined them into one species in the quantification (Figure 3 blue circles). We extracted a monomer-dimer dissociation constant of 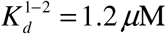 with Mg·ATPγS and 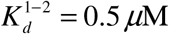 with Mg·ADP. On the other hand, the dimer-tetramer/hexamer dissociation constants were an order of magnitude smaller with Mg·ATPγS compared to Mg·ADP 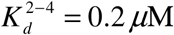 with Mg·ATPγS and 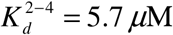 with Mg·ADP), indicating that ATPγS induces rapid formation of tetramer/hexamers from dimers.

**Figure 3.**
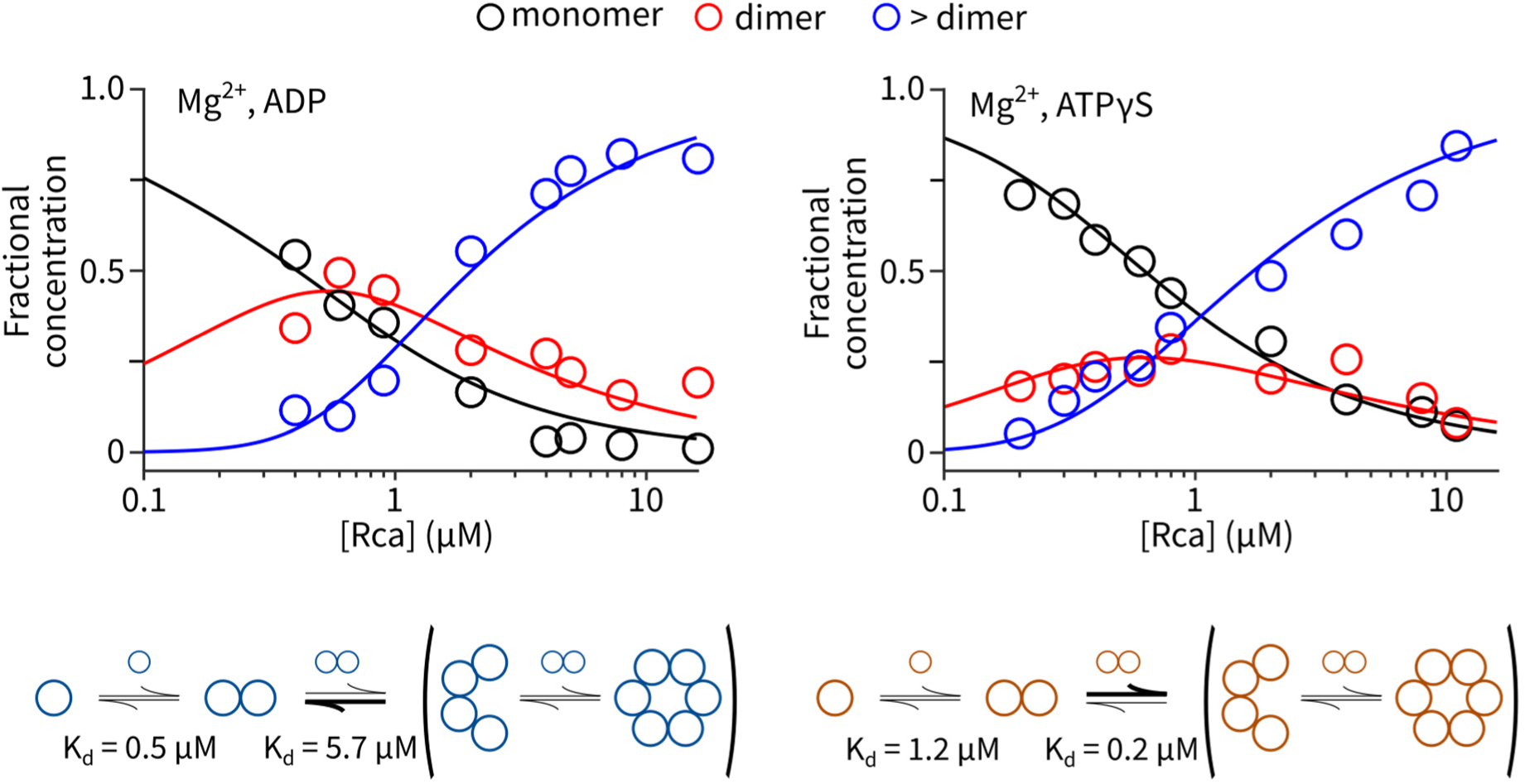
Quantifying the oligomerization equilibrium under Mg·ATPγS and Mg·ADP conditions. Solid curves are fits to the assembly model illustrated below.

### Assembly and disassembly of Nt-Rca are dynamic processes on the 1-10s timescale

Single-molecule diffusometry with the ABEL-oligo method offers the unique opportunity to observe dynamic assembly/disassembly processes that underpin an equilibrium, by measuring the time-dependent changes in hydrodynamic radius of a single protein complex. To observe these processes in real time, we optimized experimental conditions to greatly extend trapping duration of single Alexa647-labeled Nt-Rca proteins (from ~1 second to ~5-10 seconds). We indeed observed many molecules showing dynamic transitions in diffusivity with time resolution of ~150ms. Figure 4A shows two examples, both taken with Mg·ADP. In the top panel, the molecule entered the trap as a tetramer, disassembled into a monomer, and then underwent sequential assembly (monomer-dimer-tetramer/hexamer). In the bottom panel, the molecule most likely entered as a dimer and underwent multiple rounds of assembly and disassembly processes. By pooling all transitions detected under the Mg·ADP condition, we created a transition map by plotting the ending D state against the starting D state (Figure 4B). As illustrated by the transition map, the most encountered transitions are between dimers and tetramers/hexamers. The second most populated transition is between monomers and dimers. These results highlight the dynamic nature of the dimer-tetramer/hexamer pathway and further support the 1-2-4-6 assembly model we used to fit the equilibrium data. Interestingly, dynamic assembly and disassembly processes were also observed with Mg·ATPγS but with much decreased frequency, as quantified by Figure 4C. From the frequencies of the observed transitions, we estimated the rate of subunit exchange to be ~0.1s^-1^ with ATPγS and >0.2 s^-1^ with ADP. These data suggest that binding of ATPγS locks the protein into a more stable oligomer while ADP binding destabilizes the oligomers. It also suggests that some of the smearing observed in Figure 2, especially in the Mg·ADP case (for example, the trimer-like density maxima at 0.8μM), could be due to assembly/disassembly dynamics.

**Figure 4.**
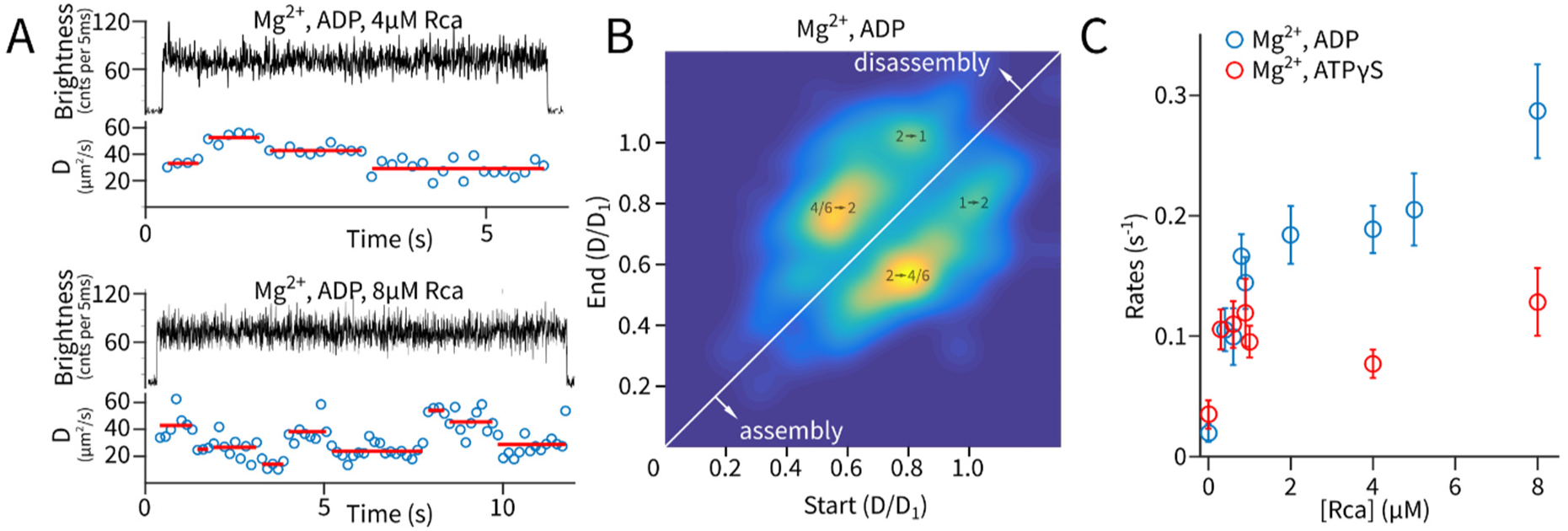
Dynamic assembly and disassembly processes observed at the single-molecule level. (A) Two example traces showing the single-molecule diffusion coefficient stepping between discrete levels. (B) Density map of all resolved transitions under the Mg·ADP condition. The first four density maxima are annotated with the interpreted transition pathways. (C) Transition rates versus protein concentration, under different nucleotide binding conditions. Error bars were estimated from the Poisson counting noise of the number of transitions.

## Discussion

We have shown that single-molecule diffusometry is a novel and powerful method to resolve protein oligomerization pathways in solution, under equilibrium conditions. It is a single-molecule (here single-complex) method, thus applicable to sub-μM concentrations and dynamically interacting samples, where traditional bulk techniques fail. The method is based on measuring a physical property (the hydrodynamic radius) of individual objects, thus providing a more direct readout compared to other single-molecule fluorescence techniques (e.g. step-wise photobleaching, FRET, etc.). The method only requires fluorescent labeling of a small fraction of the species in equilibrium and can monitor oligomerization reactions at physiological protein concentrations (>1 μM), unlimited by the concentration barrier^36^ for conventional single-molecule optical detection. Here, we demonstrated sufficient resolution to resolve monomers, dimers, tetramer/hexamers and higher oligomers. Indeed, the method will be most powerful in the small oligomer regime, where the hydrodynamic sizes of the different species differ by >20%. It should be noted that with a sufficient amount of detected photons, much smaller differences could in principle be resolved^29^. Taken together, we envision the ABEL-oligo method described here to be widely applicable to other oligomerization systems and more generally, to problems that involve biomolecular interactions.

The biological system studied here is the assembly pathway of tobacco Rubisco activase. Our data reveal that Nt-Rca assembles through a monomer-dimer-tetramer-hexamer pathway, which is consistent with the recent biophysical characterization of cotton Rca using FCS^10^,^22^ and related AAA+ family proteins^37^. We find that ATP binding (mimicked by non-hydrolyzable ATPγS in our experiments) strongly influences the assembly pathway. In particular, ATP binding promotes assembly of higher-order oligomers in the form of tetramers and hexamers. Notably, the extracted equilibrium constant for hexamer formation from dimers (K_d_^2^-^4^×K ^4^-^6^), 0.2 μM^2^, is identical to that reported for cotton β-Rca in the presence of ATPγS^22^. Quantitative analysis of the equilibrium composition and dynamic assembly/disassembly transitions of Nt Rca reveal the dimer as a critical intermediate that responds differentially to ATPγS and ADP binding. ATPγS binding drastically reduces the dimer-tetramer/hexamer dissociation constant and suppresses the rate of subunit exchange, leading to efficient and stable formation of tetramers/hexamers that dominate the equilibrium at 1-10μM protein concentration.

We hypothesize that ATP binding at the subunit-subunit interface induces a conformational change at the dimer level that “primes” the protein for stable formation of higher-order, functional complexes. Nucleotide binding could serve as a regulatory mechanism that controls the concentration of functional oligomers. We anticipate future experiments on mutants and under different ATP/ADP ratios, together with correlated activity assays, to reveal deeper structural-functional insight into this system.

## ACKNOWLEDGMENT

This work is supported in part by the Division of Chemical Sciences, Geosciences, and Biosciences, Office of Basic Energy Sciences of the US Department of Energy through Grant DE-FG02-07ER15892 (to WEM) and DE-FG02-09-ER16123 (to RMW). QW acknowledges support from the Lewis-Sigler Fellowship from Princeton University.

